# SMTdb: A comprehensive spatial meta-transcriptome resource in cancer

**DOI:** 10.1101/2025.01.22.634407

**Authors:** Weiwei Zhou, Qingyi Yang, Jiyu Guo, Si Li, Minghai Su, Feng Leng, Tingyu Rong, Jingyi Shi, Yueying Gao, Tiantongfei Jiang, Juan Xu, Yongsheng Li

## Abstract

Microbes have been found in various tumors and the research of tumor microbiome has been garnering increased attention. However, it remains challenge to investigate the microbiome in cancer at spatial resolution. The rapid advent of spatially resolved transcriptomics techniques has given rise to map transcripts at single-cell resolution in various types of cancer. Here, we constructed a comprehensive spatial meta-transcriptome resource by manually curating 203 fresh frozen (FF) slices from 20 cancers encompassing 334,253 spots and 1,908,646 cells. SMTdb (http://bio-bigdata.hrbmu.edu.cn/SMTdb/) was constructed to provide comprehensive insights into the abundance, distribution and enriched TME regions of 1218 microbiota in spatial tissue slices. SMTdb enables vast interactive data exploration of spatial distribution and expression of microbiota, host gene modules associated with certain microbiota and co-occurrence between microbiota and immune cells within tumor microenvironment. The atlas resource serves as a one-stop and time-effective platform to investigate the interactions among microbial ecosystems and hosts in cancer.

## INTRODUCTION

The tumor microenvironment (TME) is a central region of tumors with complex components. Diverse factors secreted by immune and nonimmune cells in the TME drive the inflammatory, immunosuppressive, and proangiogenic tumor internal environments (Jin and Jin 2020). Numerous studies have confirmed that the interactions between tumor cells and immune cells play a critical role in TME (Mao et al. 2021; Ma et al. 2023). In addition, emerging evidence has shown that microbes are present in various tumors and can impact cancer progression (LaCourse et al. 2021), metastasis (Bullman et al. 2017; Parhi et al. 2020; Fu et al. 2022), immune monitoring (Jin et al. 2019; Riquelme et al. 2019), and even drug resistance (Geller et al. 2017; Yu et al. 2017). Understanding the composition of the microbiota and its communication with cells in TME is vital for elucidating the molecular mechanisms of tumors.

The development of single-cell RNA sequencing (scRNA-seq) technology provides the possibility to explore the relationships between the microbiota and cells in TME. Recent study has characterized the features of intra-tumoral microbiota and revealed the most abundant bacterial orders in intrahepatic cholangiocarcinoma (Chai et al. 2023). Moreover, spatial transcriptomics (ST) technology allows exploration of the spatial distribution and co-occurrence of different cell types and microbiota in tissue slices. For example, host‒microbe interactions in oral squamous cell carcinoma and colorectal cancer have been revealed (Galeano Nino et al. 2022). However, attempts are just emerging, and there is no work to explore the spatial distribution pattern of microbiota and their interactions with immune cells in cancer.

Here, we construct a comprehensive spatial meta-transcriptome resource, SMTdb, by integrating scRNA-seq and ST data. We manually curated 203 FF slices from 20 cancer types encompassing 334,253 spots. To assess the cell type composition of spots, over 1,900,000 cells from paired or robust scRNA-seq data were collected as reference. SMTdb provides comprehensive insights into the abundance, distribution and enriched TME regions of 1218 microbiota in spatial tissue slices. Furthermore, SMTdb offers multiple analytical modules that allow users to interactively investigate the spatial distribution and expression of microbiota, host gene modules associated with certain microbiota and co-occurrence between microbiota and immune cells within TME. As the first spatial microbiota-TME analysis and data resource, SMTdb will promote microbial studies of tumor infection and the host immune response.

## RESULTS

### Spatial and single-cell transcriptomes across cancer types

To construct a comprehensive spatial meta-transcriptome resource in cancer, we manually collected FF tissue samples published through a literature search over the past five years. FF samples are considered the gold standard for spatial genomics research, as they provide higher-quality DNA and RNA than do formalin-fixed and paraffin-embedded (FFPE) samples. After strict filtering and uniformly processing, we assembled a dataset comprising 203 tissue slices from 20 cancer types, including 120 tumor slices and 83 normal slices, with a total of 334,253 spots. The tumor slices contained an average of 1,800 spots per sample, whereas the average number of normal tissue slices was 1,600 spots (Figure 1 and Supplemental Table S1).

**Figure 1:**
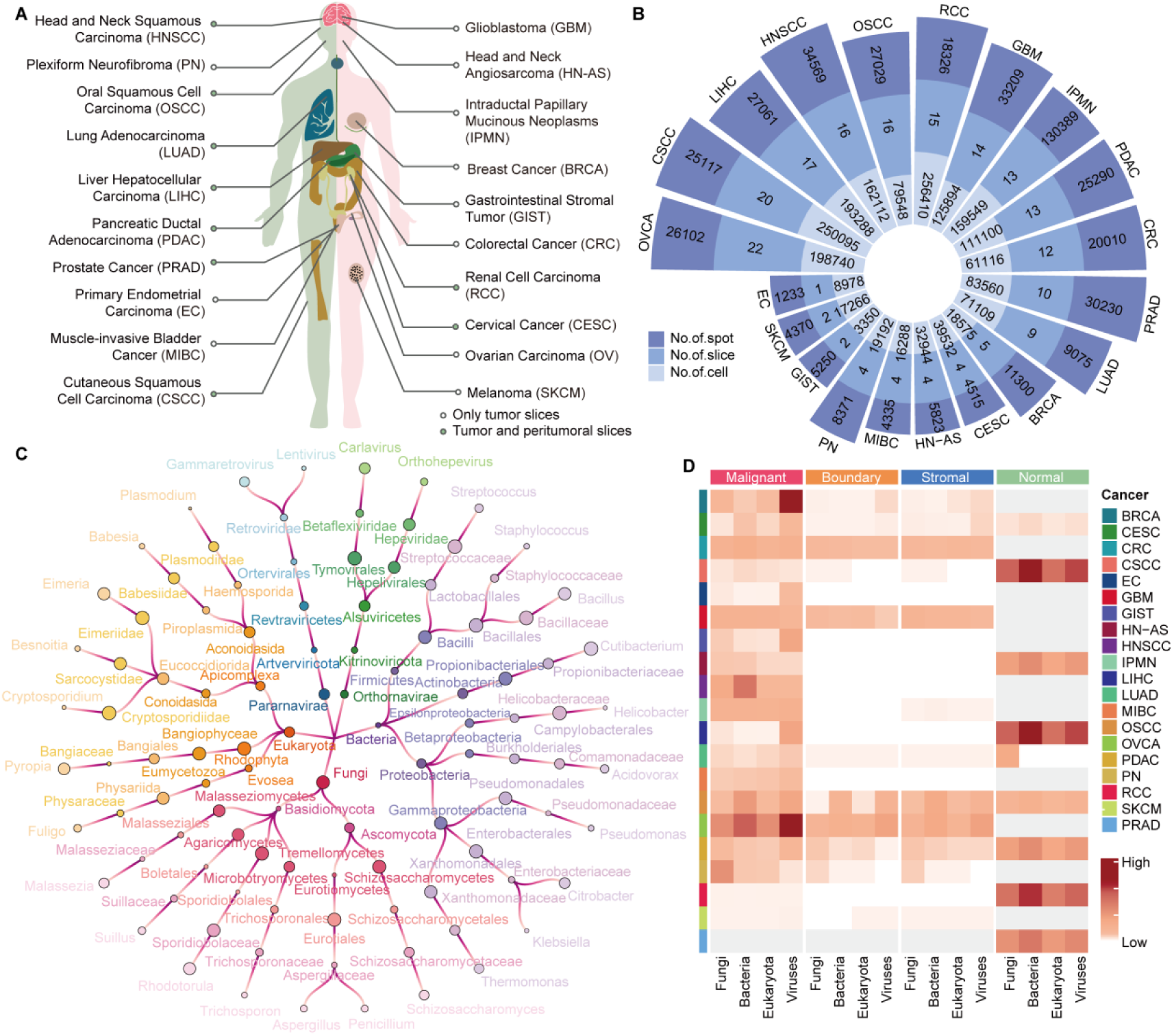
Overview of SMTdb. (A) and (B) Data collection of multi-omics. (C) Lineage map of microbiota in SMTdb. (D) Abundance of microbiota across cancer types.

Furthermore, to analyze the composition of the immune microenvironment in spatial tissue sections, we collected paired or robust scRNA-seq data and annotated cell types based on marker genes (Supplemental Table S3). After rigorous quality control and filtering, we constructed a pan-cancer single-cell atlas encompassing 1,908,646 cells across 42 cell types, including 187,114 tumor cells, 1,077,641 immune cells and 644,071 stromal cells (Figure 1 and Supplemental Table S1). Finally, using the corresponding single-cell atlas for each cancer type, we assigned cell types to each spot in tissue slices.

### Spatial meta-transcriptome in cancer

Utilizing the constructed spatial transcriptomic atlas, we applied the analysis pipeline SMT (a method for extracting microbial sequences from ST data and assigning taxonomic labels) to assess microbial abundance within each spot. We found that microbiota within the tissue slices were predominantly bacteria, fungi, and eukaryotes, along with a minor presence of viruses (Supplemental Figure 1A).

A total of 26 viruses, 309 bacterial, 391 eukaryotic, and 492 fungal species were identified (Supplemental Figure 1B). Notably, the microbiota enriched in each cancer type showed significant differences. *Citrobacter* was enriched in colorectal cancer (CRC), glioblastoma (GBM), muscle-invasive bladder cancer (MIBC), renal cell carcinoma (RCC), and ovarian carcinoma (OV), while *Acidovorax* was predominantly enriched in breast cancer (BRCA) and *Helicobacter* in gastrointestinal stromal tumor (GIST), suggesting the microbial diversity and specificity across different cancer types (Supplemental Figure 1A,D). In addition, tumor tissues exhibited a greater enrichment of microbiota implicated in cancer development and progression (Jiang et al. 2015; Stasiewicz and Karpinski 2022; Zong et al. 2023), including *Candida*, *Agaricus*, and *Malassezia*, compared to normal tissues (Supplemental Figure 1C). Our analysis constructed a comprehensive atlas of microbial distribution and abundance in spatial contexts, providing a foundation for investigating the interactions between microbiota and TME.

### User interface (UI) overview

We present SMTdb, a comprehensive data portal offering the exploration and visualization of microbial spatial distributions and their interactions with TME. To enhance usability, SMTdb provides versatile functional panels, supporting overview and data exploration.

The “Browse” page offers a slice list displaying cancer types, the number of microbiota, spots and cell types. Users can filter the table by cancer type, tissue, or microbiota of interest and select a slice to access more information via the “Detail” button (Figure 2A).

**Figure 2:**
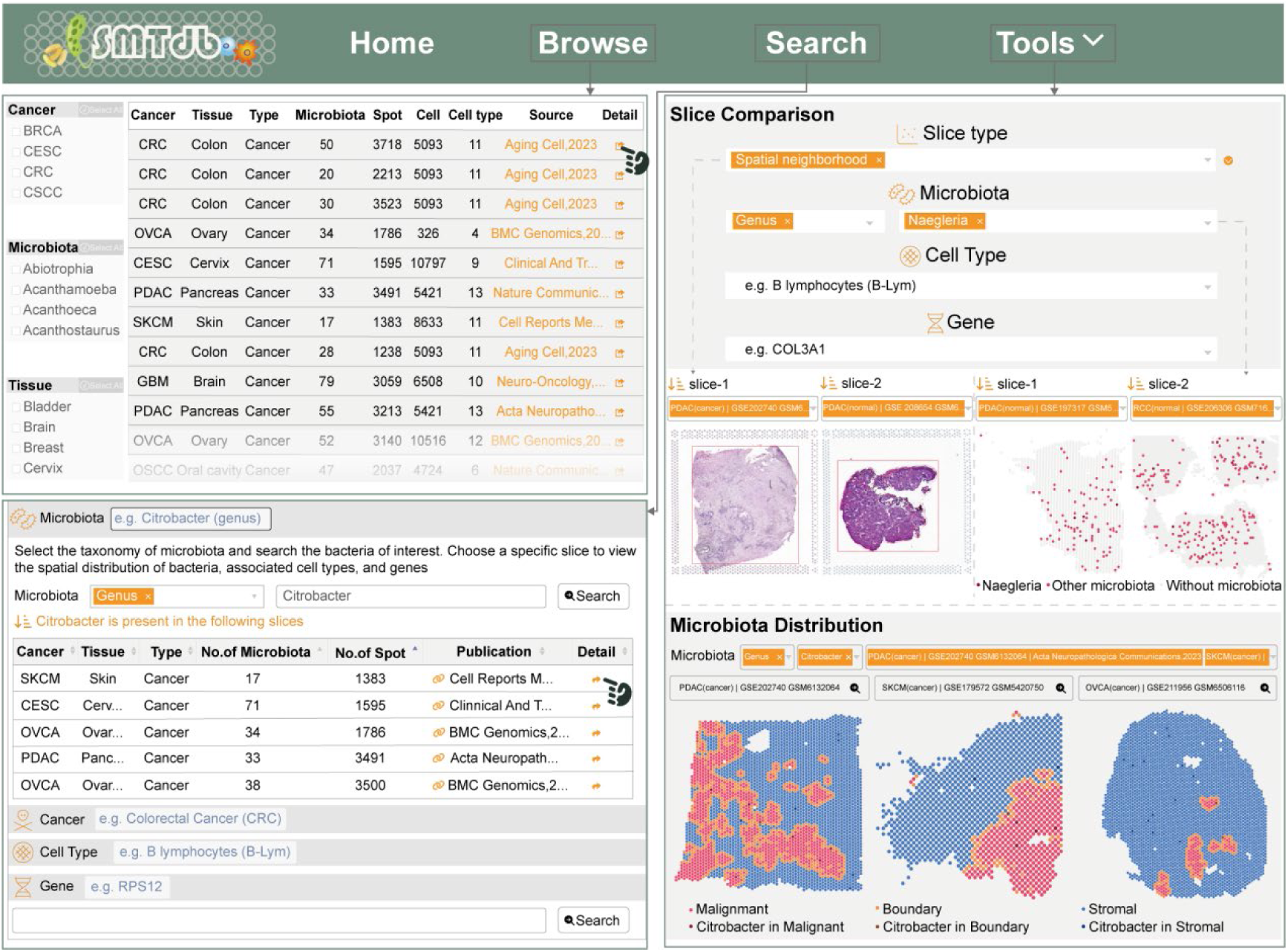
UI of SMTdb database. (A) The “Browse” page allows users to filter slices of interest using the buttons located on the left sidebar. (B) Example results generated by using “*Citrobacter*” as a key on the “Search” page. (C) On the “Slice Comparison” page, users can analyze the similarities and differences across multiple features between any two slices. (D) On the “Microbiota Distribution” page, users can visualize the spatial distribution of specific microbiota within different neighborhood regions across multiple slices.

The “Search” page offers four modes, allowing users to query specific microbiota, cancers, cell types, or genes. SMTdb filters slices to deliver tailored results. As an example, Figure 2B highlights slices containing “*Citrobacter*”.

The “Tools” page provides two interactive functions: “Slice comparison” and “Microbiota distribution”. The former aims to insight variations in the distribution of characteristics such as spatial neighborhoods, microbial distribution, cell type composition, and gene expression between two slices. Users can select any two spatial slices to explore the associations and differences between any features of interest within the slices (Figure 2C). “Microbiota distribution” is designed for rapid comparison of a selected microbiota across multiple spatial slices. Users can view the spatial distribution of the specific microbiota in slices. Moreover, quick links are provided for users to access more details related to the microbiota in the selected slice (Figure 2D).

### Slice annotation

SMTdb annotated multi-omics data for each tissue slice. First, SMTdb identified the marker genes of the transcriptome clusters. Users can browse the expression of genes within specific spatial locations. SMTdb also provided SVG for each slice, facilitating an understanding of the cellular states and functions unique to particular spatial region (Figure 3A). In addition, based on scRNA-seq, SMTdb provides the composition of cell types for each spot, enabling users to select specific spot or region for the analysis of cell localization in tissue slices (Figure 3A). Ultimately, the data related to microbiota within spatial slices were also integrated herein. Users can explore the spatial distribution and abundance of microbiota of interest across individual spot (Figure 3A).

**Figure 3:**
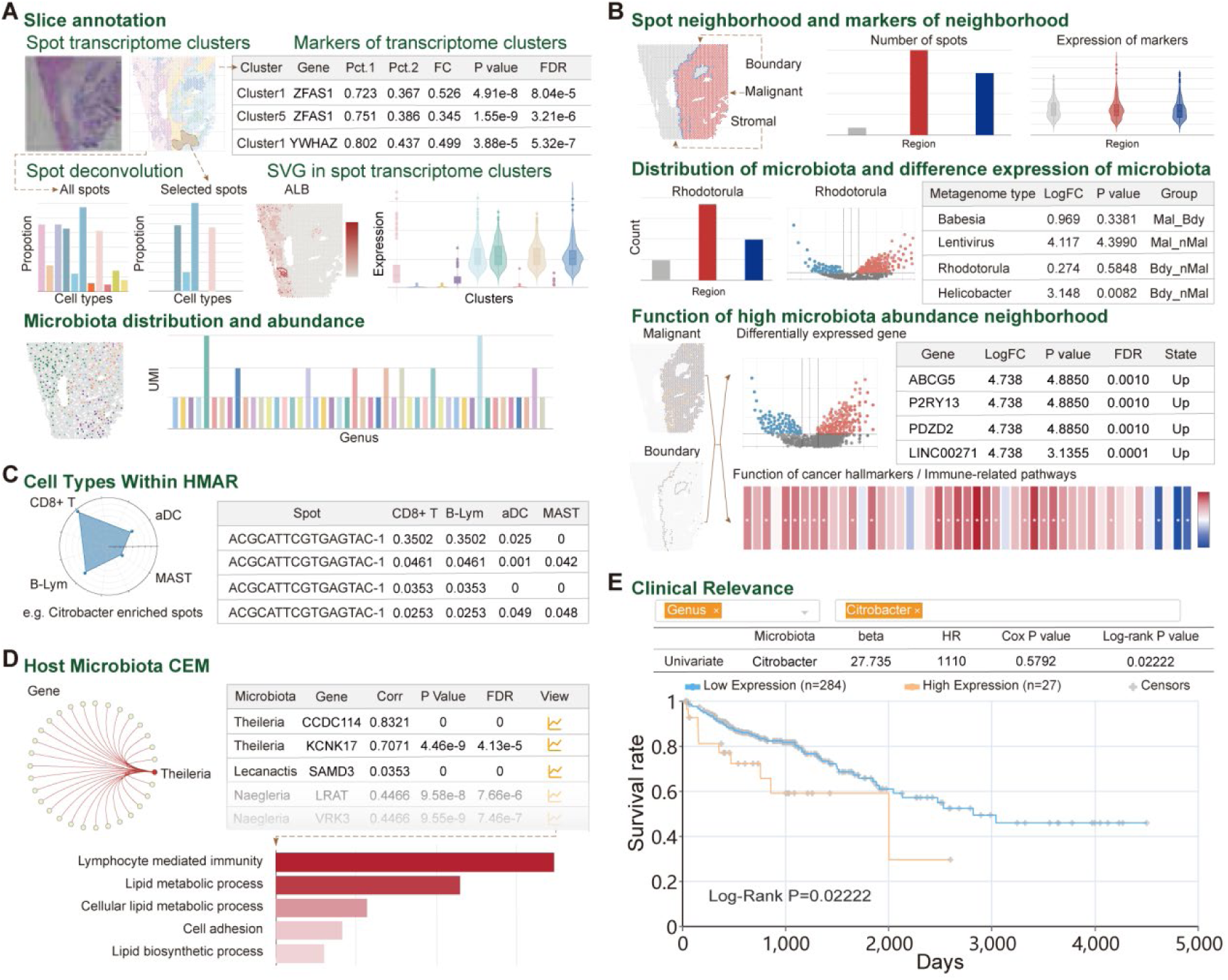
Analytic modules in SMTdb. (A) Slice annotation module offers transcriptomic clusters, marker information of clusters, deconvolution of spot and the abundance of microbiota in tissue slices. (B) Spatial neighborhood module provides the distribution of spots in spatial slice, identifies differentially regulated genes and the enriched functions by microbiota in malignant and boundary regions (cancer hallmarks for malignant spots and immune pathways for boundary spots). (C) Co-occurrence module shows immune cell types that co-occurrence with bacteria in TME across spatial context. (D) Gene modules regulated by bacteria in the host and the biological functions regulated by these modules. (E) K-M plot for CRC patients of TCGA cohort based on the abundance of Citrobacter, showing the classification of patients into high-expression and low-expression groups by the median value for analysis.

### Functions of regions with high bacterial abundance

In addition, SMTdb defines malignant and boundary regions for each tissue slice. The malignant regions consisted of cells with the most prominent oncogenic features, whereas the boundary regions encompassed the outermost layer of malignant in a solid tumor and spatially close stromal cells, which are frequently infiltrated by immune cells (Figure 3B). In the malignant region, the focus was directed towards exploring the cancer hallmarks regulated by microbiota-enriched spots, whereas in boundary regions, SMTdb emphasized the activity of immune pathways. Users can easily access these results through various types of visualizations, such as volcano plots and heatmaps (Figure 3B).

### Cell types within a high microbiota abundance region

The colonization of bacteria can lead to the proliferation of immune cells, such as T cells and B cells, influencing the development of tumors (Overacre-Delgoffe et al. 2021). SMTdb provides the composition of cell types in high microbiota abundance regions (HMAR) (Figure 3C). Users can identify the co-occurrence of different immune cells and microbiota within a spatial context, offering guidance for the comprehension of microbial functions within TME (Figure 3C).

### Host microbiota co-expression module and clinical relevance

To understand the interactions between the host and microbiota and reveal how the microbiota affects the host’s physiological and immune responses, we studied the co-expression modules of the microbiota and host genes (Figure 3D). SMTdb calculates the microbial gene co-expression modules in each slice. Users can identify the gene modules regulated by microbes and the biological functions enriched by it in the spatial slice (Figure 3D). In addition, to investigate the relationship between microbial abundance and prognosis, we performed Cox regression analysis and log-rank tests to identify outcome-associated microbial taxa at the genus level (Figure 3E).

### Case Study

Liver metastasis drives the main malignant tumor-progression events of CRC patients, whereas most studies have focused only on the cellular ecosystem of liver tissue. Recent work has revealed the remarkable roles of the microbiome in metastatic cancer. Therefore, we used an ST dataset of metastatic liver in CRC to analyze the relevance of the cellular composition and microbiota (Garbarino et al. 2023). Spatial neighborhood and spot transcriptome clusters were generated from tissue slices. Consistent with the senescent metastatic cancer cells in the original study, SMTdb identified the malignant regions and adjacent boundary regions in the same slice (Figure 4A,B). Parts of spots were annotated as epithelial signature-like metastatic cancer cells (eSMCC) in the original study, which exactly appeared in the spot transcriptome cluster 2 (Figure 4B and Supplemental Figure 2A). Similar to previous results, eSMCC-accumulated RP11 was also significantly overexpressed in cluster 2 (Figure 4C). Combined with scRNA-seq reference data, we explored the cellular composition of spatial spots. We found that monocytes dominated malignant boundary regions, which may be recruited by tumor cells (Supplemental Figure 2B). Notably, macrophages and monocytes were concentrated mainly at the interface between malignant and stromal regions, which was highly consistent with the original findings (Figure 4D and Supplemental Figure 2C). Moreover, these cells were enriched in 6, 7, and particularly 8 spot transcriptome clusters, which implied that the transcriptional clusters covered important information on the cell type distribution (Figure 4E and Supplemental Figure 2D).

**Figure 4:**
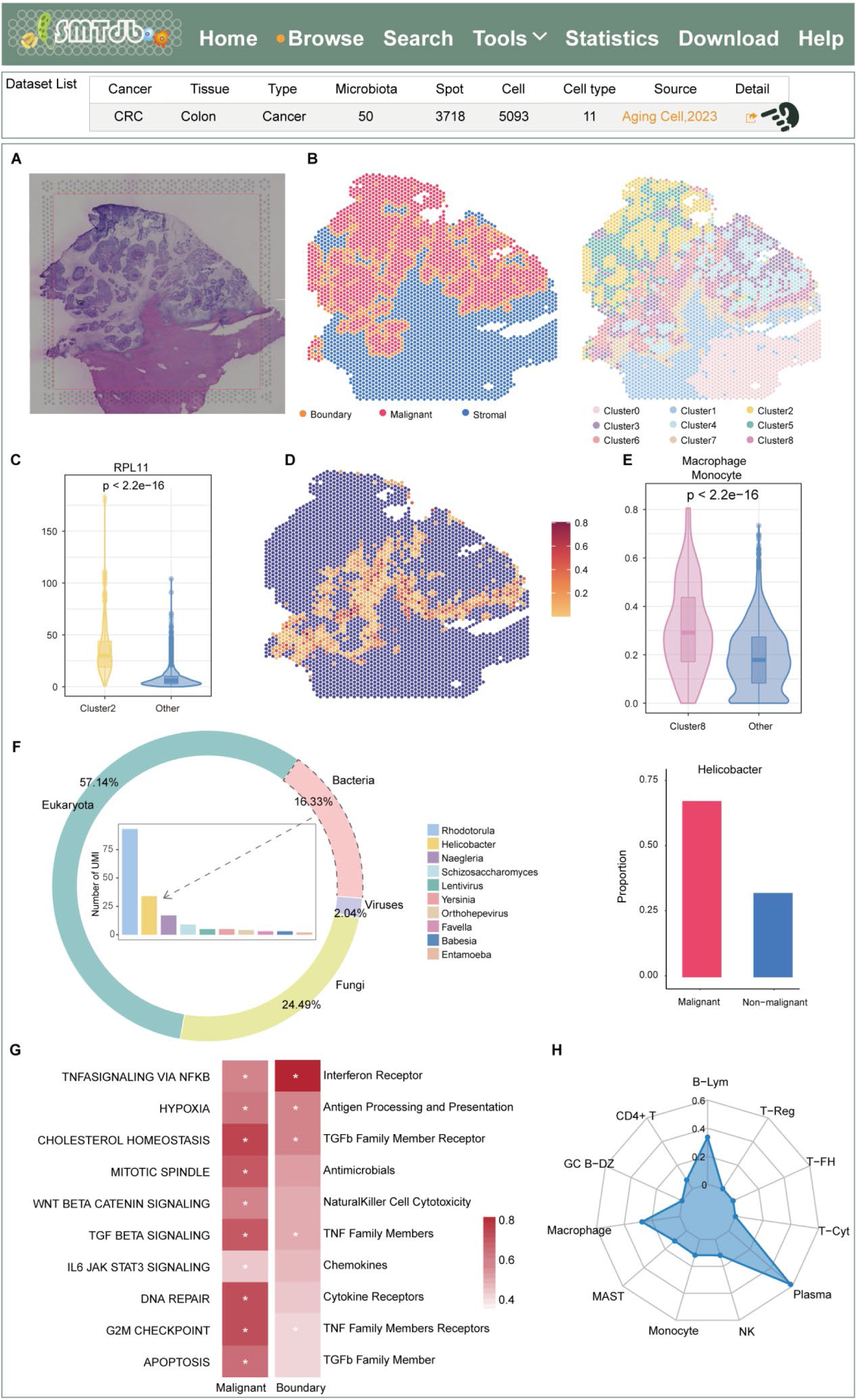
Case study of colorectal liver metastasis patients in SMTdb. (A) HE image of tissue slice. (B) Spatial neighborhood and transcriptome clusters of tissue slice. (C) Expression of RPL11 between cluster 2 and other clusters (Wilcoxon rank sum tests). (D) The distribution of macrophages and monocytes in tissue slices. (E) Proportion of macrophages and monocytes in cluster 8 compared with the other clusters. (F) The composition of microbiota within tissue slices (left) and the abundance of H.pylori in various spatial neighborhoods (right). (G) Cancer hallmarks regulated by microbiota enrich in malignant (left) and immune pathways regulated by microbiota enrich in boundary (right, * means FDR<0.05 by hypergeometric test). (H) Immune cells co-occurring with H.pylori in spatial context.

Furthermore, we performed metagenomic analysis on another CRC liver metastasis slice. Among all the spots, the second highest abundance *Helicobacter pylori* (*H.pylori*) has been found to be associated with the development of liver disease (Figure 3F)(Sumida et al. 2015; Boziki et al. 2021; Chen et al. 2023b). Additionally, as the most prevalent bacteria in the digestive system (Ralser et al. 2023), the spread of colorectal cancer cells may be the other cause of the abundance of *H.pylori* in liver tissue. As expected, *H.pylori* appeared mainly in the malignant region composed of metastatic cancer cells (Figure 4F). To investigate the function of spots enriched microbiota, we used the specifically upregulated genes in HMAR, which were identified in STMdb, for malignant and stromal regions separately Supplemental Figure 2E). Functional enrichment analysis indicated that cell cycle, DNA repair and TGF-β signal pathways were upregulated in the malignant cell region, whereas interferon receptor and antigen-related processes were activated in the adjacent boundary regions (Figure 4G). Co-occurrence analysis within STMdb indicated that *H.pylori* was closely associated with monocytes and multiple B lymphocyte subtypes in the spatial context (Figure 4H), suggesting that their interactions may play a role in the immune response (Peek et al. 2010). Besides, a previous study has elucidated that *H.pylori* induces the formation of tertiary lymphoid organs, leading to liver inflammation (Shomer et al. 2003).

In summary, based on the annotated ST datasets and analysis pipeline within STMdb, we examined the cellular composition and spatial positioning in CRC liver metastasis slices and dissected the distribution of the microbiota and the co-occurrence between the microbiota and cells, providing a possible interpretation for the liver metastasis of cancer cells and immune-microbial ecosystem in TME.

## DISCUSSION

This study integrates single-cell omics, spatial transcriptomics, and metagenomics data to construct SMTdb, the first comprehensive spatial meta-transcriptome resource for human cancer. This user-friendly platform facilitates the integration of multi-omics data, enabling users to explore the expression and spatial distribution of microbiota and cell types with clarity and precision. SMTdb addresses a critical gap in spatial metatranscriptomic data for cancer research and provides a powerful tool to investigate the interactions between microbes and TME. By combining single-cell omics, spatial transcriptomics, and metagenomics, SMTdb offers researchers a comprehensive spatial perspective on the role of microbes in tumor progression and their interactions with host cells. This enables a deeper understanding of microbial niches and functions across the TME and elucidates microbe-host interactions.

SMTdb contains 203 tissue sections from 20 different cancer types, covering 334,253 spots, 945,149 cells, and 1218 microbiota. This extensive dataset allows users to compare microbiome differences across cancer types and investigate associations between specific microbes and cancer phenotypes. SMTdb features seven analytical modules, including slice annotation (e.g., spatially co-occurring gene modules, spot deconvolution, and spatially variable genes), microbiota spatial distribution, spatial neighborhood delineation, functional analysis of microbiota-enriched regions, cell type co-occurrence with bacteria, bacterial-host gene co-expression analysis and clinical relevance of microbiota. These modules provide insights into the spatial distribution of microorganisms and cell types in tissue sections, unveiling their roles and interactions within the TME.

Despite the robust capabilities of SMTdb, it currently includes only FF slices due to sequencing technology limitations. With the continuous development of sequencing technology, in the future, we will continuously update SMTdb and expand it to other species. In summary, by enhancing the understanding of host‒microbiota interactions, SMTdb will become a valuable resource, significantly contributing to advancements in biology and medicine.

## METHODS

### Data collection

We collected raw FF ST data from the Gene Expression Omnibus (GEO) (Barrett et al. 2013) and public studies, including 120 tumor tissue slices and 83 peritumoral tissue slices from 20 cancer types (Supplemental Table S1). Furthermore, 1,908,646 cells from paired or robust scRNA-seq data (Zhou et al. 2024) were matched to each slice as reference data to assess the cell type composition of spots.

### Abundance of microbiota in spatial slices

We employed STM, a genome sequence-based pipeline, to extract microbiota (Supplemental Table S2) from ST datasets (Lyu et al. 2023). Briefly, reads that did not map to the host genome were first filtered and denoised, followed by BLAST alignment to the NCBI Nucleotide database (https://www.ncbi.nlm.nih.gov/nucleotide/), generating spatially resolved microbial abundance matrices and host gene expression profiles. Next, each spot was annotated with specific taxonomic labels if the UMI was greater than 0. The following analyses of the microbiota were based on the abundance matrix.

### Processing and clustering ST and scRNA-seq data

To obtain the spatial transcriptome expression of each spot, the raw fastq files were processed with the spaceranger tool (version 2.0, 10x Genomics) and mapped to the human reference genome (GRCh38). The Seurat R package was used for subsequent analyses (Butler et al. 2018). Only those spots coinciding with tissue slices were retained, whereas spots of low quality, which are identified by either an excessively small or large gene count per spot, were filtered. To normalize the raw counts, a strategy of regularized negative binomial regression, specifically the SC transform (Hafemeister and Satija 2019), was adopted. Principal Component Analysis (PCA) was then applied to achieve dimensionality reduction. The Louvain algorithm (resolution = 0.8) was used to generate transcriptome clusters. To identify the marker genes associated with each cluster, the ‘findAllMarkers’ algorithm was executed, adopting the parameters ‘min.pct = 0.25, logfc.threshold = 0.25’. To identify spatially variable genes (SVGs), we used the function ‘FindSpatiallyVariableFeatures’ in Seurat to measure the complex expression patterns of genes (SVGs; FDR < 0.05 and I > 0).

For the analysis of all the scRNA-seq datasets, a uniform analytical pipeline was adopted utilizing the Seurat tool. Cells exhibiting high mitochondrial expression and total unique molecular identifier (UMI) counts were filtered to ensure data quality. Furthermore, normalization of the raw UMI counts was conducted via the SC transform. Following this preprocessing step, highly variable genes were identified to facilitate the distinction of cellular heterogeneity. PCA was then used to enable robust clustering of the cells.

### Cell type annotation for scRNA-seq data

A total of 363 canonical marker genes were manually recorded for 42 cell types from our previous work (Supplemental Table S3) (Jiang et al. 2023). The markers were used as reference atlas for a fully automated and ultra-fast cell type identification method, ScType, to assign cell types to each cell (Ianevski et al. 2022). InferCNV (version 1.2.1) (Patel et al. 2014) was employed to identify malignant cells based on copy number variations (CNVs), with immune cells serving as reference cells.

### Identification of differentially expressed genes

To analyze the impact of bacterial presence on gene expression in tumors, we divided the malignant, boundary, and stromal spots into two groups based on whether bacteria were present in the spots and conducted differential expression analysis. The Wilcoxon rank-sum test was utilized to assess the statistical significance of differentially expressed genes, and the false discovery rate (FDR) method was applied for p-value correction. Genes whose FDR value was less than 0.05 and whose fold change (FC) value was greater than 1.5 were considered significantly upregulated.

### Identification of spatial neighborhood

To decode the complex spatial microenvironment of the tumor, we employed Cottrazm (Xun et al. 2023) to map the microenvironment at the tumor boundary. The SME normalization algorithm from the stLearn (Pham et al. 2023) package is used to adjust gene expression based on the spot image matrix, resulting in a morphologically adjusted gene expression matrix (Morph), and spatial spots are clustered via the KNN algorithm from the Seurat package. Immune-related gene signatures were scored in the Morph matrix to define a normal tissue expression score (NormalScore) for each spot. Cottrazm selects a reference based on the highest median NormalScore within the cluster. InferCNV (Patel et al. 2014) was employed to assess CNV levels for the remaining spots.

Hierarchical clustering was performed to categorize spatial spots into clusters to distinguish malignant spots and stromal spots. The CNV scores of each spot were incorporated into the Seurat object, and spots with high median CNV scores were initially defined as core malignant spots. Cottrazm calculated centroids for malignant and normal clusters and determined the proximity of each spot to these centroids. Spots were labeled as malignant or boundary based on their relative distances. The method arranges spatial spots on hexagonal lattices and defines neighboring spots using Manhattan distances.

### Identification of spatial co-expression modules

To identify spatial co-expression netwok modules, hdWGCNA (Morabito et al. 2023) was used to analyze spatial transcriptome data. We used the MetaspotsByGroups() function to establish metaspots separately for each Seurat cluster, and the SetDatExpr() function was applied to construct the metaspot expression matrix. The soft power was tested via TestSoftPowers(), and the optimal threshold was determined. he ConstructNetwork() function was employed to construct gene co-expression network modules.

### Functional analysis

To predict the function of microbiota-enriched regions, we identified DEGs by the Wilcoxon rank sum test (FDR < 0.05 and FC > 1.5) for both microbiota-enriched and non-microbiota enriched regions separately. To assess the functions of bacteria, we estimated single-sample gene set enrichment analysis (ssGSEA) scores (Hanzelmann et al. 2013) for immune-related signatures (Li et al. 2020) and cancer hallmarks (Liberzon et al. 2015). Additionally, a hypergeometric test was performed to ascertain the functions significantly upregulated in the microbiota-enriched regions.

### Spatial correlation between host genes and microbiota

We assessed microbiota-gene co-expression modules in slices based on their expression at each spatial spot. For any pair of bacteria 𝑖 and gene 𝑗, we first obtained the probabilities 𝐸_𝑖_ = [𝑒𝑖_1_, 𝑒𝑖_2_, . . . , 𝑒𝑖_𝑛_] and 𝐸_𝑗_ = [𝑒𝑗_1_, 𝑒𝑗_2_, . . . , 𝑒𝑗_𝑛_] that were observed across *n* spots, which represented the expression of bacteria and genes in spots. Next, the Spearman correlation coefficient *R* was calculated based on the following formula:

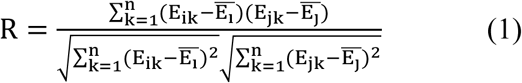

where 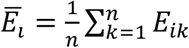 and 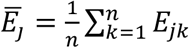 and n was the number of spots in the given slice.

For each bacteria, all positively correlated genes (R>0.1) were considered as co-expression modules. Furthermore, we investigated the functions that were influenced by the module, using hypergeometric test to Gene Ontology biological process (GO-BP).

### Survival analysis

To evaluate the association between microbial abundance and clinical prognosis we collected clinical data from The Cancer Genome Atlas (TCGA) (Tomczak et al. 2015), and the microbial abundance of TCGA samples was collected from BIC (Chen et al. 2023a). We used univariate and multivariate Cox regression analyses. Additionally, patients were stratified into high- and low-abundance groups according to the median expression of microbes in each cancer type, and survival differences between the two groups were assessed using the Kaplan-Meier (KM) method.

## DATA ACCESS

All data in the SMTdb can be downloaded from the download page. The source code has been made available on GitHub and can be accessed through the following link: https://github.com/ComputationalEpigeneticsLab/SMTdb.

## ACKNOWLEDGMENTS

This work was supported by the National Natural Science Foundation of China (32322020, 32170676, 32060152); Natural Science Foundation of Heilongjiang Province (Key Program) (ZD2023C007). We thank the National Natural Science Foundation of China. We would like to thank Natural Science Foundation of Heilongjiang Province (Key Program). We would also like to thank the member of the lab for helpful discussion and suggestion.

## AUTHOR CONTRIBUTIONS

**Conceptualization:** Juan Xu, Yongsheng Li, Tiantongfei Jiang.

**Data curatrion:** Weiwei Zhou, Qingyi Yang, Jiyu Guo.

**Formal analysis:** Weiwei Zhou, Qingyi Yang, Jiyu Guo.

**Investigation:** Weiwei Zhou,Jiyu Guo, Qingyi Yang, Si Li.

**Methodology:** Weiwei Zhou, Qingyi Yang, Jiyu Guo, Minghai Su, Jingyi Shi, Yueying Gao, Feng Leng, Tingyu Rong.

**Software:** Weiwei Zhou, Qingyi Yang, Jiyu Guo.

**Validation:** Weiwei Zhou, Qingyi Yang, Jiyu Guo., Si Li.

**Visualization:** Weiwei Zhou, Qingyi Yang, Jiyu Guo.

**Writing – original draft:** Weiwei Zhou, Yongsheng Li, Juan Xu, Qingyi Yang, Tiantongfei Jiang.

**Writing – review & editing:** Weiwei Zhou, Yongsheng Li, Juan Xu, Qingyi Yang, Tiantongfei Jiang.

